# Binding during sequence learning does not alter cortical representations of individual actions

**DOI:** 10.1101/255794

**Authors:** Patrick Beukema, Jörn Diedrichsen, Timothy Verstynen

**Affiliations:** Center for Neuroscience, University of Pittsburgh, Pittsburgh, Pennsylvania, 15260; Center for the Neural Basis of Cognition, University of Pittsburgh & Carnegie Mellon University, Pittsburgh, Pennsylvania, 15213; Brain and Mind Institute, Departments of Statistics & Computer Science, University of Western Ontario; Department of Psychology, Carnegie Mellon University, Pittsburgh, Pennsylvania, 15213

## Abstract

As a movement sequence is learned, serially ordered actions get bound together into sets in order to reduce computational complexity during planning and execution. Here we examined how the binding of serial actions alters the cortical representations of individual movements. Across five weeks of practice, healthy human subjects learned either a complex 32-item sequence of finger movements (Trained group, N=9) or randomly ordered actions (Control group, N=9). After five weeks of training, responses during sequence production in the Trained group were correlated, consistent with being bound together under a common command. These behavioral changes, however, did not coincide with plasticity in the multivariate representations of individual finger movements, assessed using fMRI, at any level of the cortical motor hierarchy. This suggests that the representations of individual actions remain stable, even as the execution of those same actions become bound together in the context of producing a well learned sequence.

## INTRODUCTION

Being able to combine simple movements into coordinated sets of actions is critical to many everyday skills, such as typing on the computer or driving a manual transmission car (Lashley, 1951). Over the course of evolution the brain has solved this sequencing problem multiple times, resulting in many interacting algorithms that facilitate the consolidation of complex skills (for review see Beukema and Verstynen 2018). One of these algorithms is the process of set building, also known as chunking or binding (Verwey 1996). Binding serial actions into sets improves computational efficiency during the production of complex actions by representing multiple movements under a single selection command (Ramkumar et. al, 2016). To illustrate this process consider the graphical model presented in Figure 1A-B. On each trial, the manual response to a visual cue occurs through a hierarchical system of perception, selection (e.g., key), and motor planning (e.g., finger movement), that are all represented as latent states with their own independent sources of noise. In this example, the serial order of cues across trials follows a deterministic sequential order. Prior to training, each response is planned independently of the preceding trial. Once the order of cues is learned, the brain can consolidate the selection process so that a set of motor plans is represented under a single selection state. This selection state is triggered by the presentation of the first stimulus in the series, after which subsequent motor commands are cued by the internal state, rather than by the visual cues. This results in faster production of responses to items within a set, as well as a correlation in responses within bound sets due to their shared upstream command (Figure 1C; Verstynen et al., 2012; Acuna et al., 2014; Lynch et al., 2017).

**Figure 1:**
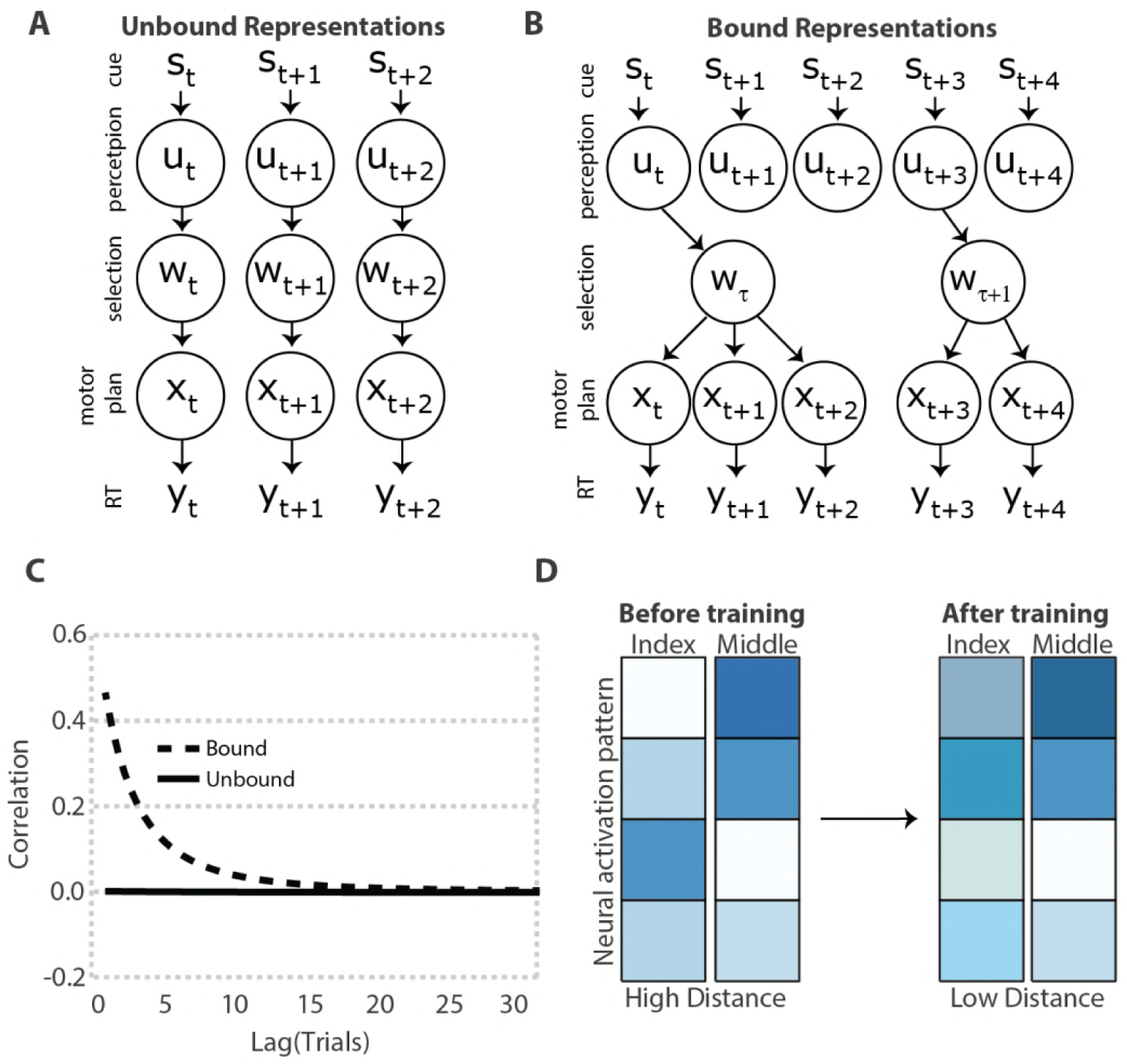
The process of response binding **A**. One each trial, (t), a visual stimulus (s) triggers an appropriate finger response (y), in this case reflecting a response time (RT). In the case of unbound actions, the visual perception (u), selection (w), and motor planning (x) processes are all represented as latent states that operate independently across trials. **B**. With training, the intermediary process of selection binds multiple motor plans together as a set. Each set of actions, τ, is triggered by the visual stimulus of the first item in the set. Subsequent actions are then internally triggered, rather than relying on external visual cues. This example shows two bound sets, a three item set followed by a two item set. **C**. The autocorrelation function of response times for bound actions (cyan) should exhibit a significant correlation across trials, while unbound actions (black) should not exhibit a temporal autocorrelation. **D**. A schematic of four hypothetical voxels in cortical sensory motor networks during the execution of either the index or middling finger, with darker colors reflecting stronger movement-evoked responses. Before training, each finger representation is associated with a unique neural activation pattern. After training, the representations of bound finger movements share more activation and the neural activation patterns are more similar.

Many forms of non-sequential motor learning rely on the reorganization of movement representations in motor networks (Nudo et. al. 1996). Therefore, it is possible that action binding during sequence learning also alters internal motor representations of individual movements; however, this effect has been largely unexplored. Recent advances in representational analysis now allow precise quantification of the relationship between the cortical activity patterns for single finger movements using fMRI (Diedrichsen and Kriegeskorte, 2017). Using this approach, previous work has shown that the structure of individual fingers in primary motor cortex is organized in a way that is consistent with their co-articulation during natural hand movements (Ejaz et. al., 2015). Furthermore, artificial manipulations of pairwise finger correlations, by physically yoking two fingers together, alters the distance between finger representations in primary somatosensory cortex (Kolasinski et. al. 2016). This suggests that elementary sensorimotor representations may be plastic and subject to changes over time and that multivariate pattern analyses on fMRI data are sensitive enough to detect these changes.

If individual actions are bound under a common motor command, then the internal representations of those actions, at some level of the motor hierarchy, should change over time in these areas that binding occurs. The naïve version of this model is that if two movements are executed repeatedly in a close temporal sequence, then the activation of one finger movement may already pre-activate the following movement. In the extreme, this model makes the prediction that two fingers that are regularly paired together in everyday actions will become enslaved together over time, thereby reducing behavioral flexibility (Lashley, 1951). It is therefore more likely that the process of binding alters the representation of contextually cued actions in upstream regions linked to more abstract response selection (Diedrichsen and Kornysheva, 2015), such as the dorsal premotor cortex (PMd) or motor regions along the intraparietal sulcus. Wherever this binding process happens, the multivariate activity pattern for the two bound movements should become more similar in that region (Figure 1D).

Here we tested the plasticity of individual action representations using a combination of behavioral analysis and event-related fMRI. Binding was measured behaviorally by looking at the degree of correlation between successive behavioral responses after training on a unimanual 32-item sequence. We also measured the population-level representations of visually-cued single finger movements in the cortex both before and after five weeks of training on the complex sequence. The simple plasticity hypothesis states that binding of serial actions after consolidation of a motor sequence should make the neural representations for those actions more similar, thereby decreasing the representational distance between them.

## RESULTS

### Learning-related changes in behavior

Participants executed sequences in a serial reaction time task (Nissen and Bullemer, 1987) on a laptop keyboard, in which each finger press was cued by a unique fractal image (Figure 2A).Subjects learned the mapping of cue to finger press prior to the first day of training. After a key press, the response time, measured as the elapsed time between cue onset and key press, was recorded and the next cue was presented following a 250 ms interval. Each day, participants were tested on trial blocks of random sequences (blocks 1,2,6) or trial blocks composed of a specific 32-element sequence (blocks 3,4,5,7). An additional control group received the same amount of training as the trained group – but here all blocks consisted of random sequences.

**Figure 2:**
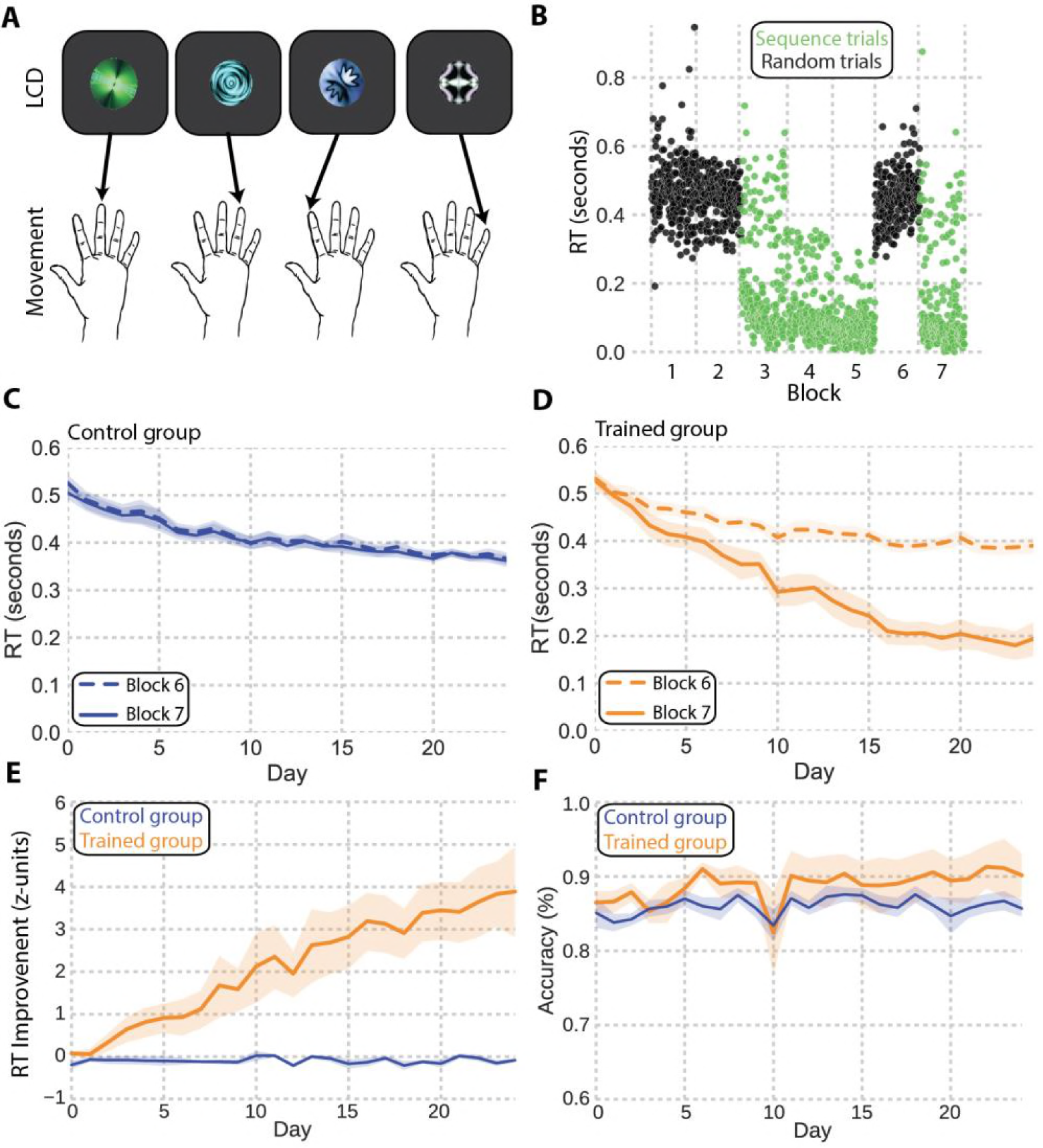
Task design and behavioral performance. **A**. Participants practiced a serial reaction time task in which each finger movement was prompted by a unique cue. **B**. Representative reaction time plot from Day 12. Each dot represents the response time on one trial. **C**. Reaction times for the Control group for random trials on blocks 6 and 7. **D**. Reaction times for the Trained group for the random trials (block 6) and sequence trials (block 7). **E**. Mean z-scored reaction times as a function of day for the Control group (blue) and Trained group (peach). **F**. Mean accuracy (correct trials/total trials) in the final trial block, as a function of day, for the Control group (blue) and Trained (peach) group. Shaded regions in panels C-F show standard error.

To assess how training impacted performance, we compared the evolution of response times and accuracy across days for the Trained and Control groups. Figure 2B illustrates all trial-wise responses during a single day for a subject in the Trained group. While responses during random trial blocks (black dots) remained relatively constant, the response times during sequence trial blocks (green dots) get steadily faster with training. The last two trial blocks were used to probe learning across time. On average both the Control (dashed line, Figure 2C) and Trained subjects (dashed line, Figure 2D) exhibited a general improvement in response speeds during the final random trial block (block 6). This general across-session speeding of responses during a trial block with random sequences likely reflects the improved learning of the cue-response mapping across days. During the final sequence block (block 7), however, sequence-specific responses in the Trained group also decreased rapidly across training days. Repeated measures ANOVA indicated a significant block × time effect: F(368,23) = 15.366, p = 7.93 × 10^-41^, with average response times dropping just below 200ms at the end of training (solid line, Figure 2D). As expected, this effect was not observed in the Control group, F(368,23) = 0.77, p = 0.76, where the final trial block did not contain an embedded sequence (solid line, Figure 2C). In order to capture sequence-specific changes in response speed, we normalized the mean response time for the final trial block (sequence in Trained group, random in Control group) by the mean and variance of response times during trial block 6 (random in both groups; see Methods). This analysis depicts a steady improvement in sequence specific response times across the 5 weeks for the Trained group, with sequence block responses approximately 4 standard deviations faster than the random trial blocks at the end of training (Figure 2E). Repeated measures ANOVA indicated a significant group by time effect, F(368,23) = 12.79, p = 1.67 × 10^-34^. Unlike response speed, average accuracy during the final trial block gradually rose at a steady rate for both groups, saturating at around 90% for the Trained group and 85% for the Control group, with no significant between group differences F(368,23) = 0.36, p = 0.99.

There are several ways that responses could get faster during the sequence blocks (see Beukema and Verstynen, 2018). The binding hypothesis (Figure 1B), however, makes the specific prediction that serially successive actions that are bound under a shared motor plan should exhibit a correlation in their responses over time, as a consequence of arising from a common, high-level motor plan (Figure 1C). For an index of binding, we used the autocorrelation of RTs during the last trial block for both groups (Verstynen et. al. 2012). Figure 3 shows the autocorrelation functions for early (Day 1), middle (Day 12), and late (Day 24) stages of practice for the Trained (Figure 3A) and Control (Figure 3B) groups separately. Participants in the Trained group showed no evidence of an autocorrelation in their RTs at Day 1; however, by the middle of training a pronounced autocorrelation of temporally adjacent responses emerged that remained steady by the last day of practice. As expected, the Control group did not show any autocorrelation structure in RTs at any point during training, indicative of a lack of binding across responses.

**Figure 3:**
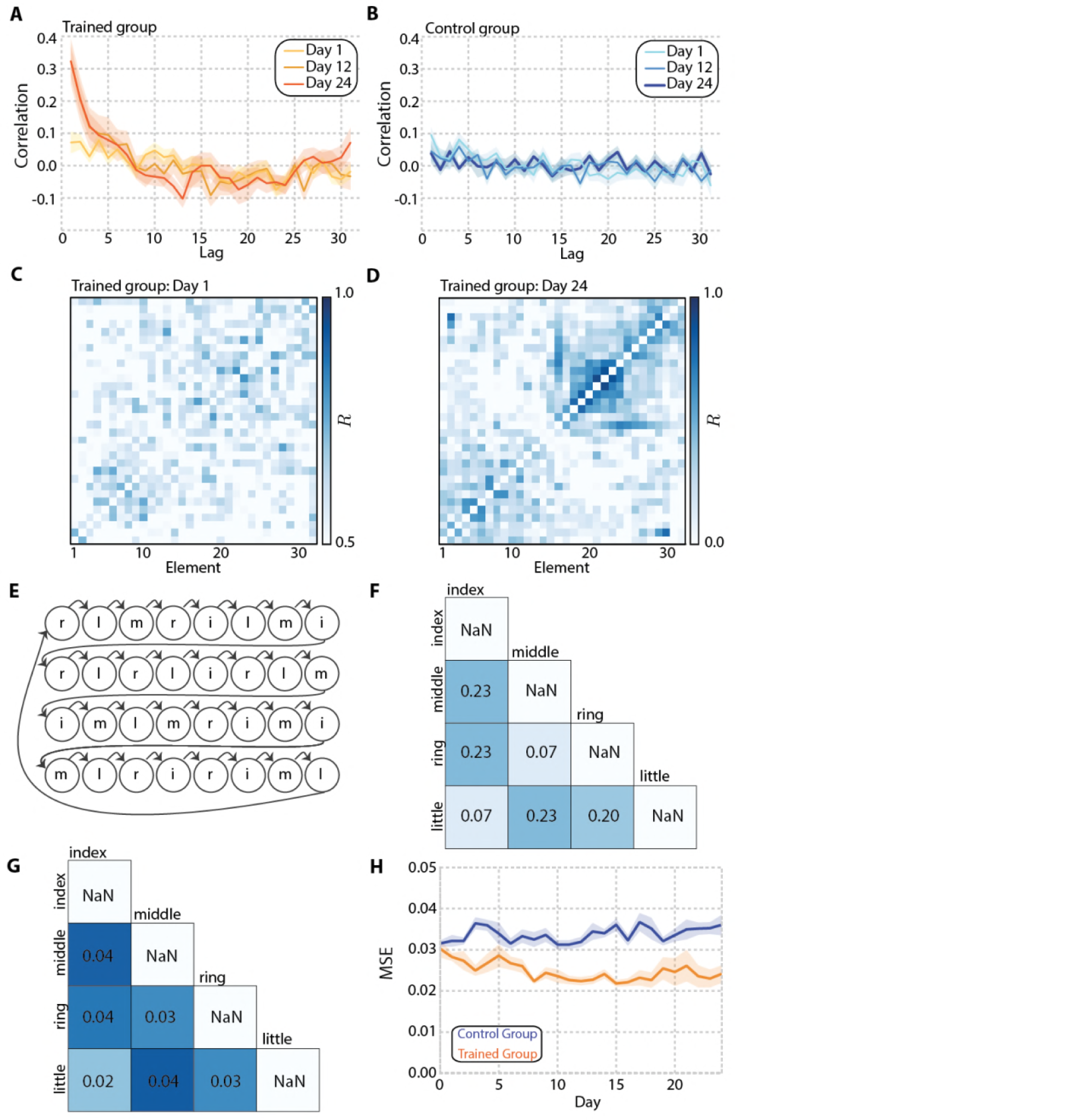
Binding in behavioral responses. **A,B** Mean autocorrelation function for lags 1-31 during early (day 1), middle (day 12) and late training (day 24) for the Control group (A) and Trained group (B). Shaded regions show standard error of the mean. **C,D**. Average correlation between each element in the sequence during the final trial block for the Trained group, during Day 1 (C) and Day 24 (D). **E**. The 32 element sequence showing frequency of each finger transition (i-index, m-middle, r-ring, l-little) **F**. Pairwise frequencies between each finger **G**. Average observed correlations between fingers at the end of training collapsed across subjects. **H**. The MSE between the pairwise frequencies (panel F) and observed correlation matrix computed separately for each subject. Smaller numbers indicate increased similarity to the expected pairwise frequencies (panel F). Shaded regions show standard error.

We next examined the RT correlations of responses to each item in the 32-item sequence on each cycle of the sequence production. Before practice, this 32×32 correlation matrix does not show much structure, with all items approximately equally uncorrelated (Figure 3C). After training, a clear structure in the correlations emerged, with local clusters of correlated responses found along the diagonal of the matrix (Figure 3D). If these clusters of correlated responses in the sequence reflect the inter-finger transition probabilities (Figure 3E), then the pairing frequency of individual fingers should determine the degree of similarity between finger responses. The similarity between the observed correlations and expected correlations based on the pairwise frequencies (Figure 3F) was computed using the mean squared error (MSE). The mean observed correlation matrix across all subjects on the final day of training is shown in Figure 3G. There was increased similarity between the observed and expected correlations across days (Figure 3H) in the Trained group, but the structure in the Control group remained unchanged, resulting in a significant group by time interaction, F(368,23) = 1.90, *p* = 0.0079.

### Stable motor representations after training

In order to directly measure multivariate cortical representations of the individual cued movements, we used a rapid-event-related fMRI design consisting of presentations of each cued finger press followed by a period of fixation (Figure 4A). A regions of interest (ROI) analysis was performed on the cortical motor network including primary motor cortex, M1; primary somatosensory cortex, S1; dorsal premotor cortex, PMd; ventral premotor cortex, PMv; supplementary motor area, SMA; and the superior parietal lobule, SPL. These regions were anatomically localized using Brodmann areas extracted from Freesurfer (see Methods). These regions are shown on the group average surface (Figure 4C). In each of the cortical motor ROIs, we quantified the activity pattern related to each cued finger movement and then calculated a cross-validated Mahalanobis distance (crossnobis) between the activity patterns for each cued finger pair. If two cued fingers generate the same cortical activity patterns, then the corresponding distance between them will be 0. However, if two finger movements consistently generate dissimilar finger patterns, then the corresponding distance will be positive (Figure 4B). Cross-validation allows us to test the value of the distance estimates directly against zero (see Diedrichsen and Kriegeskorte 2017, Walther et. al. 2016, Diedrichsen et. al. 2016). The distances between every possible pair of fingers is summarized in a representational dissimilarity matrix (RDM) for each ROI (Figure 4D).

**Figure 4:**
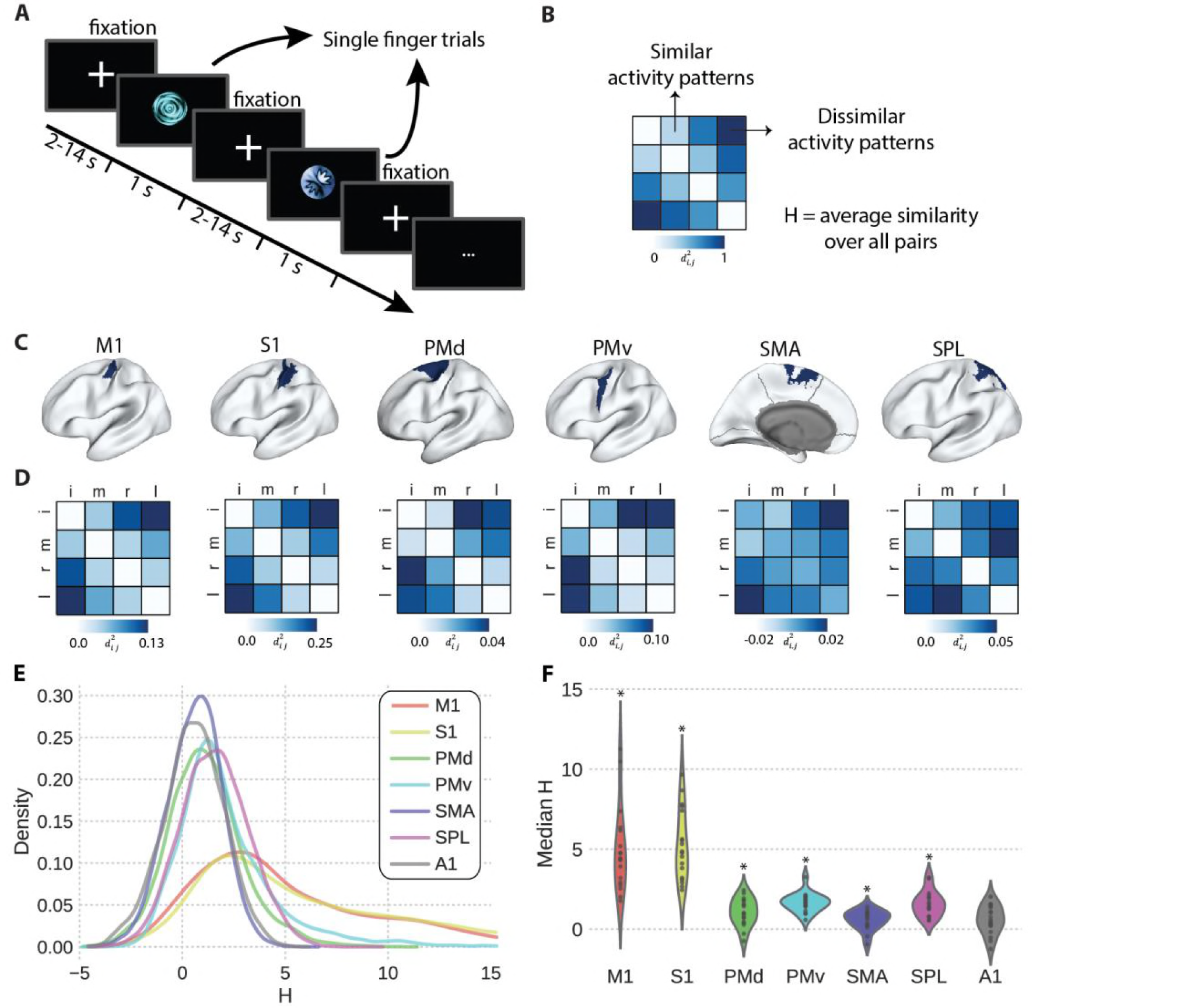
Multivariate activity patterns during cued finger movements. **A**. fMRI task schematic. Participants executed single finger movements on the button glove following a variable period of fixation. The cue-finger mapping was identical to that used during the training. **B**. Example of a representational dissimilarity matrix showing similar finger patterns that result in small distances and dissimilar finger patterns that result in large distances. The average crossnobis similarity (H) was used as a test statistic for assessing decoding in each ROI and for assessing representational plasticity. **C**. Regions of interest masks overlaid in blue on the group average surface. **D**. Average representational dissimilarity matrices for each region. Each colored square within the RDM indicates the distance between those two fingers (i=index, m=middle, r=ring, l=little) **E**. Average kernel density estimates of H across all subjects and all voxels in primary motor cortex (M1), primary somatosensory cortex (S1), premotor dorsal cortex (PMd), premotor ventral cortex (PMv), superior parietal lobule (SPL), supplementary motor area (SMA), and primary auditory cortex (A1). **F**. Violin plots show the distributions of median H values across subjects. Black circles inside plots show individual data. Asterisks indicate significance at α = 0.05 after correcting for multiple comparisons (Bonferonni).

The representational structure we observed in primary motor and primary somatosensory cortex qualitatively matches previous reports (Ejaz et. al. 2015), such that the index finger is furthest from the little finger, while the middle and ring fingers are close together. This pattern of representational distances is also similar to what is observed the other cortical motor regions, although the overall between effector distances are smaller in these premotor regions (Figure 4D). To confirm that each region has reliably different representations for the fingers, we computed the average cross-validated pairwise distance between all finger movements (Figure 4B see Methods). Average H scores greater than 0 indicate above-chance encoding (Diedrichsen and Kriegeskorte 2017). Figure 4E shows the mean H distribution computed from the surface-based searchlight (see Methods) across all voxels as kernel density estimates averaged across subjects (one distribution per subject for each ROI). In a control region, primary auditory cortex (A1), distances are symmetrically distributed about zero, indicating that one would not be able to reliably decode the cued-finger movement from this region. In sensorimotor regions, the distances were positively skewed indicative of cued-finger movement encoding. In order to estimate the reliability of this encoding across subjects, we extracted the median distance value from each distribution for each subject and ROI. A one-sample t-test on those median values (one median per subject), after adjusting for multiple comparisons using a Bonferonni correction, found significant separation of cued finger representations (i.e., positive average distances) in the cortical sensorimotor areas, but not the A1 control region (Table 1). Thus, consistent with previous studies (Ejaz et. al. 2015), the patterns of activity in the motor network can reliably discriminate individual effectors.

**Table 1:** Column “mean” indicates the average median H value in each region. T-statistics computed from one-sided t-test.* indicates significance after correcting for multiple comparisons.

| Region | Mean | t(17) | p-value | 95% CI |
| --- | --- | --- | --- | --- |
| M1 | 4.92 | 7.91 | *$4.23 \times 10^{-7}$ | 3.61, 6.23 |
| S1 | 5.23 | 10.13 | *$1.28 \times 10^{-8}$ | 4.14, 6.32 |
| PMd | 1.07 | 5.30 | *$5.83 \times 10^{-5}$ | 0.64, 1.49 |
| PMv | 1.66 | 12.06 | *$9.21 \times 10^{-10}$ | 1.37, 1.95 |
| SMA | 0.57 | 4.08 | *$7.75 \times 10^{-4}$ | 0.28, 0.87 |
| SPL | 1.57 | 8.44 | *$1.74 \times 10^{-4}$ | 1.18, 1.97 |
| A1 | 0.57 | 2.78 | 0.013 | 0.14, 1.01 |

To determine whether the emergence of binding in the behavioral responses coincides with alterations of these representational distances of individual cued actions, we measured how average distances changed for each cortical motor ROI before and after training. More specifically, if binding results in the representations of frequently paired actions becoming more similar (Figure 1D), then distances between frequently paired movements would decrease after practice only in the Trained group. When looking at all pairwise distances (Figure 5A) we were unable to find a reliable influence of sequence training on the average pattern distances in any cortical motor region (Figure 5B). In most areas, the distances decreased only marginally for both Trained and Control groups together, but the finger patterns remained largely separable, with patterns exhibiting a high degree of stability. Across all regions, we failed to detect a reliable interaction between group and time that would be indicative of a training effect in representational distances (all p>0.26, full statistics reported in Table 2). In order to evaluate the evidence in support of the null hypothesis that the interaction is not present, we conducted a JZS Bayes Factor (BF) ANOVA with uniform prior across all models (JASP Team, 2017, jasp-stats.org) and found evidence in support of the null model that training does not influence distances. The BF’s ranged from 0.33-0.36 (Table 2), which can be considered positive evidence in support of the Null hypothesis (Kass & Raftery, 1995).

**Figure 5:**
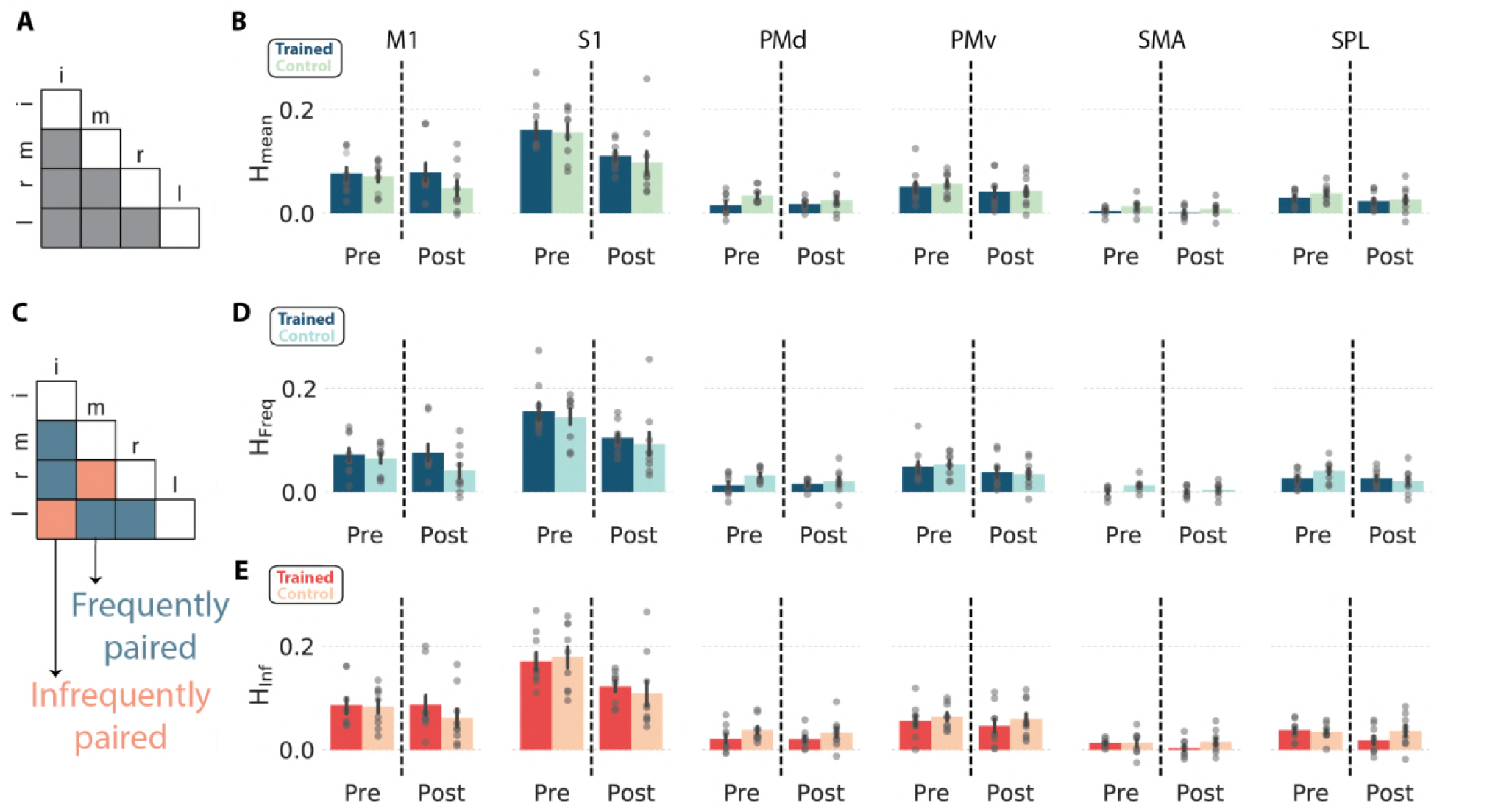
Stable representational distances after training. **A** Pairwise finger distances included in overall distance analysis. **B**. Bar plots show mean ROI H values in the pre‐ and post-training scans separately for each group. Error bars show standard error. Gray circles are individual data points. **C**. Finger pair frequencies were asymmetrically distributed in the trained sequence (see Figure 3F). Some finger pairs, e.g. index and little were infrequently paired, whereas other finger pairs e.g. index and middle were frequently paired. **D,E**. Bar plots show mean H for frequent pairs B (panel D) and infrequent pairs (panel E) in the Pre and Post scans separately for each group. Error bars show standard error. Gray circles are individual data points. No comparison was found to be statistically significant at α=0.05.

**Table 2:** F-statistics and p-values for testing significance of interaction effect (group x time) from repeated measures ANOVA for mean distances. Inclusion Bayes Factor (BF) is the ratio of the posterior over the prior probability of the model including the interaction term. BF<1 provide evidence that interaction effect is not present.

| Region | F(1,16) | p-value | Inclusion BF |
| --- | --- | --- | --- |
| M1 | 1.35 | 0.26 | 0.348 |
| S1 | 0.086 | 0.77 | 0.349 |
| PMd | 0.74 | 0.40 | 0.349 |
| PMv | 0.066 | 0.80 | 0.367 |
| SMA | .042 | 0.84 | 0.333 |
| SPL | 0.34 | 0.57 | 0.35 |

Of course, looking at changes in overall representational distances my not be sensitive enough to pick up changes in the representational distances of only a few finger pairs. The simple plasticity model we proposed in the Introduction predicts that the greatest plasticity should be observed in the finger pairs most often executed together in the sequence. If the distances decreased for the more frequently paired effectors, but increased for the less frequently paired effectors this may result in a net change for the overall average distance near 0. To explore this possibility, we re-analyzed the distance changes by looking at the frequently and infrequently occurring finger pairs in the sequence structure itself (Figure 3F). Based on the pairing frequencies, we identified four frequent finger pairs and two infrequent pairs (Figure 5C). However, much like the overall distance patterns, we were unable to resolve focal changes in representational distances in either of the most frequently (Figure 5D) or infrequently (Figure 5E) paired effectors. Across all regions, two way repeated measures ANOVA indicated no significant group-by-time interaction for either frequently paired (all p > 0.26, full statistics provided in Table 3) or infrequently paired fingers (all p > 0.13, full statistics provided in Table 4). The Bayesian ANOVA revealed anecdotal evidence in favor of the null hypothesis for both the frequently (BFs: 0.68-0.89, Table 3) and infrequently (BFs: 0.53-0.60, Table 4) paired fingers.

**Table 3:**
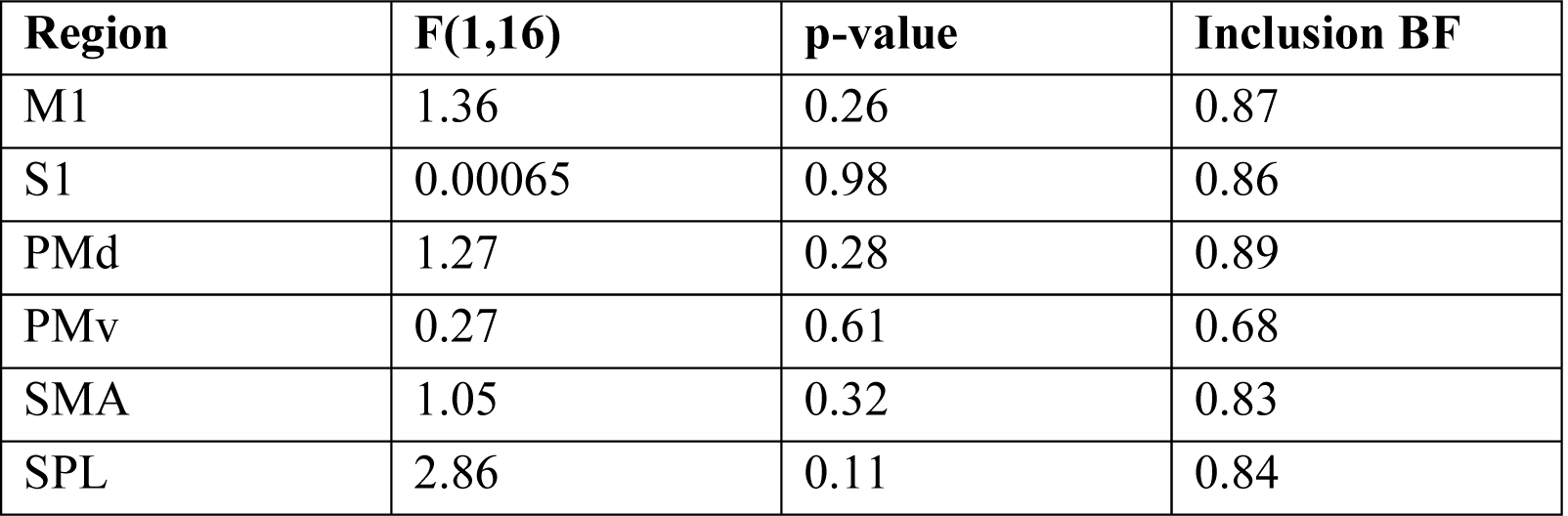
F statistics and p-values for testing significance of interaction effect (group x time) from repeated measures ANOVA for frequently paired fingers. Inclusion Bayes Factor (BF) is the ratio of the posterior over the prior probability of the model including the interaction term. BF<1 provide evidence that interaction effect is not present.

**Table 4:** F statistics and p-values for testing significance of interaction effect (group x time) from repeated measures ANOVA for the infrequently paired fingers. Inclusion Bayes Factor (BF) is the ratio of the posterior over the prior probability of the model including the interaction term. BF<1 provide evidence that interaction effect is not present.

| Region | F(1,16) | p-value | Inclusion BF |
| --- | --- | --- | --- |
| M1 | 1.16 | 0.30 | 0.54 |
| S1 | 0.69 | 0.42 | 0.60 |
| PMd | 0.070 | 0.80 | 0.54 |
| PMv | 0.083 | 0.78 | 0.53 |
| SMA | 1.05 | 0.32 | 0.56 |
| SPL | 2.53 | 0.13 | 0.54 |

## DISCUSSION

Here we examined whether the binding of serial actions during long-term sequence learning alters the cortical representations of individual cue-response pairings. We found that during sequence production, temporally adjacent responses develop a high degree of correlation in their response speeds, consistent with participants binding multiple responses together under a unified command so as to reduce computational complexity (see also: Verstynen et. al. 2012, Ramkumar et. al. 2016, Lynch et. al. 2017). Using a multivariate pattern analysis approach, based on the cross-validated Mahalanobis estimator, we also replicated previous studies showing that cortical motor areas reliably distinguish between activation patterns of individually cued finger responses (Ejaz et. al. 2015). We were, however, unable to find evidence for learning-related changes in this representational structure of cued finger responses in any of the cortical regions tested. Taken together, these findings show that the process of binding actions into chunked sets during long-term skill learning does not impact the representation of individual cued actions, suggesting that binding relies on changing more complex levels of representation beyond individual movements.

At first glance, the absence of plasticity in population level representations of individual actions that we observed appears to be incompatible with previous reports of plasticity in sensorimotor cortex. Kolansinki and colleagues (2016) found that the representational distances of individual fingers shifted in S1 after physically yoking two fingers together for a period of 24 hours. In their study, the sensory representations of the two yoked fingers remained spatially and temporally identical, however the unyoked fingers altered their distances, suggesting a possible compensatory effect in the sensory representations themselves. In contrast, our task relied on training associations between temporally independent movements in a specific context. It is possible that, had we trained on chord-like movements, where multiple fingers are simultaneously engaged (Verstynen et. al. 2005), for a longer period of time, we might have observed similar changes in cortical sensorimotor representations, a hypothesis that is left open to future studies.

Alternatively, there is a strong rationale for why single effector representations would remain stable in cortical sensorimotor networks, particularly motor execution areas like M1, after long-term sequence learning. First, binding responses at the execution level may be a maladaptive strategy for maintaining a flexible movement repertoire (Lashley, 1951). For example, if index finger movements were consistently bound with middle finger movements because a single daily task required them to work together in sequential fashion, then they might exhibit a prepotent response in inappropriate contexts. In order to maximize flexibility, it would be beneficial for the movements to be bound at a more abstract motor planning stage, upstream from execution processes. Second, practice may involve refining the control of execution-level representations without necessarily impacting the representations themselves. This would suggest that the process of binding during the consolidation of complex movement sequences is dependent on plasticity mechanisms at hierarchically higher level of processing (Wong et. al. 2015).

Of course, it is possible that there is plasticity in the representations of individual sensorimotor effectors during long-term sequence learning, but limitations in our experimental design may preclude identifying those changes. First, while the duration of training we used was longer than many sequence learning experiments in humans, five weeks may still not be enough time to lead to measurable representational changes in primary motor cortex. This concern is tempered by the fact that we were able to show strong evidence of action binding in the behavioral responses. Second, we could not look at finger representations in the striatum, where there is some evidence for binding (Jin et. al. 2014, Wymbs et. al. 2012) as the voxel sizes in this study were too large to examine those representational structures. Future studies at a higher MRI field strength (e.g., 7T) may afford a better spatial resolution for picking up plasticity of sensorimotor representation in the striatum.

Despite these limitations, our experiment clearly shows that five weeks of training on a complex unimanual sequence task does not alter the sensorimotor representations of individual effectors despite clear evidence of binding in the motoric actions. This suggests that execution level representations remain stable during learning and that proficiency is likely controlled by a higher level within the motor hierarchy.

## MATERIALS AND METHODS

### Participants

Eighteen right-handed participants (6 female, mean age: 26 years) were recruited locally from Carnegie Mellon University (CMU) and the University of Pittsburgh. Two authors (PB and TV) were included in the sample. All participants provided informed consent and were financially compensated for their time. All experimental protocols were approved by the Institutional review board at CMU.

### Serial reaction time task

Participants were trained for 25 nonconsecutive days on a variant of the serial reaction time task (Nissen and Bullemer, 1987). All experimental procedures were performed on a laptop running Ubuntu 14.04. At the beginning of each training session, participants were instructed to place their right hand over the ”h” (index),”j” (middle), ”k” (ring), and ”l” (pinky) key. Each trial consisted of a presentation of one of four unique fractal cues appearing on a black background. Each cue was uniquely mapped to one of four keys on the keyboard (Figure 2A). The trial ended either when the participant executed a response or once the maximum response window expired (see below), depending on which event happened first. After a trial termination, the next cue was presented after a 250 ms inter-trial interval. Each trial block consisted of 256 trials and was followed by a rest period where the mean response time (RT) and accuracy for that block was provided to the participant. On each training day, participants completed 1792 trials, separated into 7 trial blocks. RT was calculated as the delay between stimulus presentation and a key press. Stimulus presentation and recording was controlled with custom written software in Python using the open source Psychopy package (Peirce, 2007). The software used for training is available on GitHub (CoAxLab, n.d.).

Prior to the first session, subjects were assigned to either a Trained group (n=9) or a Control group (n=9). For participants in the Trained group, trial blocks were separated into two types: blocks of pseudo randomly ordered cues (Random; blocks 1,2,6) or blocks of deterministically ordered cues following an embedded 32-element sequence (Sequence; blocks 3,4,5,7). Figure 2B shows the blockwise structure for a single subject in the Trained group. Trials during the Random blocks were constrained such that repeated presentations of the same cue were excluded. This was done so that Random trial blocks would appear more similar to the Sequence trial blocks. The 32 element sequence presented on Sequence blocks consisted of the following key presses: 3-4-2-3-1-4-2-1-3-4-3-4-1-3-4-2-1-2-4-2-3-1-2-1-2-4-3-1-3-1-2-4 using the mapping (1-index finger, 2-middle finger, 3-ring finger, 4-little finger). Each Sequence block began in a random position of the sequence. For the first 2 blocks, the response threshold for each trial was set to 1000 ms. To encourage faster responses, the response window of blocks 3-5 was adaptively controlled such that the response window on one trial block was the mean plus one standard deviation of the RTs from the previous trial block. If that value fell below 200 ms or if the accuracy on the preceding block was less than 75%, the threshold was reset to 1000 ms. The threshold was removed for the final probe blocks (6 and 7) so that participants could move as quickly as they chose. For the Control group, the procedure was nearly identical to the Trained group, with the exception that all 7 blocks consisted of pseudorandomly ordered trials, i.e. there was no exposure to Sequence blocks.

### Analysis of training data

Python code and source data to generate all figures is publicly available on GitHub (https://github.com/CoAxLab/binding_manuscript). All behavioral analysis during training focused on responses during the last two trial blocks (probe blocks) when no adaptive response window was applied: Random and Sequence conditions for the Trained group, Random and Random conditions for the Control group. Differences in response time (RT) and accuracy (percent correct responses) were measured as the difference in the means between the last two blocks, normalized by the standard deviation of values in trial block 6, i.e., z-scored difference in performance (Verstynen et al., 2012). In the Trained group this reflects the sequence specific change in performance on each day. Since 3 subjects completed 24/25 days of training, average group visualizations are presented for day 24 so as to evaluate the same state of learning for all subjects.

Binding was measured by computing the autocorrelation of the series of RTs within each probe trial block. The first 32 trials were excluded to remove the exponential decay as it distorts the autocorrelation analysis (Verstynen et al., 2012). The linear trend was then removed by regression and the residuals were used to calculate the autocorrelation function for lags 1 through 31, following the same procedure as described in (Verstynen et al., 2012; Lynch et al., 2017).

Since the autocorrelation function measures general associations across all sequential lags, it is not sensitive to specific associations between individual elements, and therefore cannotbe used to measure binding between specific finger pairs. Therefore, we conducted a secondary analysis on the same data but examined pairwise correlations between each distinct element (1-32) in the sequence. Average correlations, ordered by sequence element, are shown in Figure 3C-D. Binding between successive elements is reflected by increases in correlations distances before compared to after training.

To measure how much the correlation between finger responses matches the statistical structure of the trained sequence, we collapsed the elementwise correlation matrices by finger identity (index, middle, ring, pinky), forming 4×4 observed correlation matrices. To measure the similarity of the observed binding structure to the expected binding structure, we computed the mean squared error between the finger pairing frequencies of the sequence and observed correlations. This gives a normalized similarity measure for how well the pattern of correlations in the behavioral responses matches the pairwise similarities of the trained sequence.

### Imaging acquisition

Before and after training, all participants were scanned at the Scientific and Brain Research Center at Carnegie Mellon University on a Siemens Verio 3T magnet fitted with a 32-channel head coil. High-resolution T1-weighted anatomical images were collected for visualization and surface reconstruction (MPRAGE, 1 mm isotropic, 176 slices). A fieldmap with dual echo-time images (TR: 746 ms, TE1: 5.00 ms, TE2: 7.46 ms, 66 slices, 2 mm isotropic) was acquired to correct for fieldmap inhomogeneities. For the functional imaging sessions, we acquired 241 T2* weighted echo-planar imaging volumes (2 mm isotropic, TR: 2000ms, TE: 30.3 ms, MB factor: 3, 66 slices, A >> P, FoV: 192 mm, interleaved ascending order, flip angle: 79 deg, matrix size: 96×96×66, slice thickness: 2.00 mm). For the finger mapping task, we collected a total of 6 runs resulting in 1446 volumes. Functional images were oriented so as to maximize coverage of the entire cortex and cerebellum.

### Neuroimaging tasks

We collected a set of finger mapping runs to estimate the activation patterns evoked by performing each distinct cue-response pair in isolation (i.e. not embedded within a sequence). The same stimuli from the behavioral experiments were projected on an MR-compatible LCD screen mounted at the rear of the scanner. Participants could see this screen through a mirror mounted on the head coil. Responses were recorded on a five-key MR compatible response glove (PST Inc.) placed under the right hand. Each effector (e.g., individual cue-response pairing) was presented in isolation on each trial with no structured order between trials. Thus, the paradigm only measured responses to individual cued movements, not the sequence itself. Each trial type was repeated 12 times per run totaling 72 trials per session. Subjects were instructed to press the cued key several times following stimulus presentation until the cue disappeared from the screen (1 second). Between runs, subjects were given the option to take several minutes of rest.

### Imaging Analysis

Functional imaging data were analyzed using SPM8 (http://www.fil.ion.ucl.ac.uk/spm/) and custom Matlab and Python functions. Raw functional EPI images were realigned to the first volume. No slice time correction was applied. These realigned images were then corrected for field distortions using the field maps. All analyses were performed in native functional space. Structural T1 images were used to reconstruct the pial and white surfaces using Freesurfer (Fischl, 2012). All custom code is publicly available (CoAxLab, n.d.).

All analyses of task-related responses were performed using a region of interest (ROI) approach. Anatomical ROIs were defined separately for each subject, using the surface based Brodmann areas extracted from Freesurfer (Fischl et al., 2008) following similar conventions as described in (Wiestler and Diedrichsen, 2013). The hand voxels of the primary motor cortex (M1) were defined as the surface nodes with the highest probability of belonging to Brodmann area (BA) 4, 1 cm above and below the hand knob (Yousry et al., 1997). S1 was defined as the nodes in BA1 BA2, BA3a, or BA3b, 1 cm above and below the hand knob. Premotor cortex was defined as the nodes belonging to BA6 medial (PMv) or lateral (PMd) to the medial frontal gyrus. Supplementary motor area (SMA) was defined as the voxels in BA6 along the medial wall. The Freesurfer atlas was used to define the superior parietal gyrus, as it is not defined by a unique Brodmann area. As a control ROI, we extracted the voxels belonging to primary auditory cortex as this region would not be expected to exhibit any significant decoding of the visually-cued finger patterns. Each surface based ROI was projected back into native functional space.

Analysis for effector representations was performed using representational similarity analysis (RSA) using the crossnobis estimator (Kriegeskorte et al., 2008; Nili et al., 2014; Walther et al., 2015). A GLM with regressors for each effector was fit for each mapping run, along with the six head motion regressors (x, y, z, pitch, yaw, roll). Omissions and incorrect key presses were regressed out of the model. Raw time series were orthogonalized by eigenvector decomposition and projected into the principal component space to minimize model bias in the decoding. To estimate the differences between finger patterns, we used a cross-validated Mahalanobis distance between prewhitened regression coefficients for each effector (Atsushi et. al., 2017, Walther et al., 2015). The cross-validated Mahalanobis distance has the advantage over other distance measures in that it is unbiased, since noise is orthogonalized across runs, resulting in an expected distance of 0 if a voxel or region does not reliably distinguish two finger patterns (Ejaz et al., 2015). The estimated distance (*d̂*) between the patterns (*u*) of two fingers (i,j) was averaged across every pair (m,l) of runs (M) resulting in (6 choose 2) = 15 folds using the following equation:

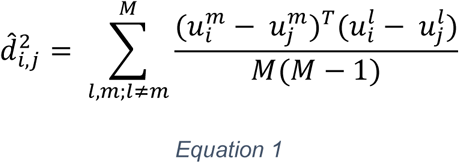

The pairwise distances between each of the fingers are summarized in a representational dissimilarity matrix (Figure 4B). To test for encoding and plasticity within each voxel or ROI, we extracted the average distance between each pair of fingers pattern (K=4) using the following equation:

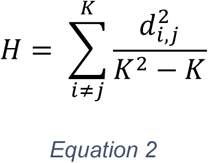

To examine the extent of finger representations across all of cortex, we conducted a surface-based searchlight (Oosterhof et al., 2011), assigning every surface node an H value based on the local (p=160) patterns surrounding an approximately 10 mm radius. Values for the number of voxels (p) and radius were chosen based on previous studies (Yokoi et. al. 2017). This searchlight approach enabled us to examine the entire H distribution across all voxels in each of the ROIs to confirm that each region reliably discriminated individual effectors. Due to the observed positive skew, we extracted the median H for all regions across all subjects and conducted a one sample t-test against 0, in order to establish whether a region reliably decoded the single finger movement representations. For tests of plasticity, changes in representational distances were compared using the patterns across the top 150 voxels from each ROI, rank-ordered by average distance, similar to the number of voxels used in previous studies (Wiestler and Diedrichsen, 2013), because representational geometries are highly sensitive to the number of voxels that make up a pattern (Oosterhof et al., 2011). We computed H separately for the pre and post training sessions and each ROI. A repeated measures ANOVA was used to examine the influence of training on distances in each ROI. Bayesian repeated measures ANOVA with a JZS prior over all models was used to determine the inclusion Bayes Factor to measure the extent to which the data supported inclusion of the interaction effect (JASP Team, 2017, jasp-stats.org). The guidelines in (Kass and Raftery, 1998) were used to interpret the weight of the evidence in support of the null hypothesis.

## Acknowledgements

The authors would like to thank Kyle Dunovan and Kevin Jarbo for helpful comments on this manuscript. Patrick Beukema received support from the Multimodal Neuroimaging Training Program NIH T90 DA022761. This research was sponsored by the Pennsylvania Department of Health Formula Award SAP4100062201 and National Science Foundation CAREER Award 1351748.

